# Contrasting segregation patterns among endogenous retroviruses across the koala population

**DOI:** 10.1101/2023.11.16.567356

**Authors:** Mette Lillie, Mats Pettersson, Patric Jern

**Author notes:** Correspondence: Mette Lillie and Patric Jern.

## Abstract

Koalas (*Phascolarctos cinereus*) are subject to retroviral infections by the koala retrovirus (KoRV), resulting in accumulation of heritable endogenous KoRV in the koala germline. Like other vertebrates, koalas have experienced a history of retroviral epidemics leaving marks as endogenous retroviruses (ERVs) in their genomes; another recently identified ERV lineage, named *phaCin-β*, shows a pattern of recent, possibly current, activity with high insertional polymorphism in the koala population that predates the establishment of endogenous KoRV. We investigate geographic patterns of retrovirus activity, focusing on three distinct ERV lineages of increasing estimated ages from KoRV to *phaCin-β* and to *phaCin-β-like*, using a whole-genome sequencing dataset of 430 koalas produced by the Koala Genome Survey. We find thousands of ERV loci across the koala population and identify contrasting patterns of polymorphism. In the youngest established ERV-lineage (KoRV), we identify thousands of integrations among northern individuals. In the *phaCin-β* ERV lineage, which is slightly older yet with overlapping time of activity with KoRV, we observe hundreds of integrations in northern individuals. In southern koalas, however, *phaCin-β* loci are found at higher frequencies possibly reflecting a bottleneck effect. Overall, the findings suggest high ERV burden in koalas that represent historic retrovirus-host interactions. Importantly, the ERV catalogue supplies improved markers for population conservation genetics of this endangered species.

## Introduction

Retroviruses are a diverse group of RNA viruses that require conversion of their RNA genome to DNA, which integrates irreversibly as a provirus into the host’s nuclear DNA. Infection normally occurs in somatic cells, where the provirus shares the cell’s fate and will thus not persist over time. If the germline is infected, however, the provirus can be transmitted to offspring as a heritable endogenous retrovirus (ERV). Over the evolutionary time-scale, ERVs have accumulated in contemporary genomes and offer a glimpse into the past retrovirus infection history^1,2^.

Koalas (*Phascolarctos cinereus*), an iconic feature of Australian wildlife, have attracted considerable attention. In addition to their cultural significance and conservation concerns, they are also hosts to the koala retrovirus, KoRV^3^, which occurs across a large range of the eastern koala distribution. KoRV has wide-ranging disease outcomes, including cancers and opportunistic infections^4,5^, and is also observed in the koala germline as endogenous KoRV (enKoRV)^6-8^.

We recently reported that, in addition to the *gammaretroviral* enKoRV, the koala germline also contains recent accumulation of a *betaretroviral* ERV lineage, named *phaCin-β*, which began infiltrating the koala genome between 0.7 and 1.5 million years ago^9^. Here, we investigate the insertional polymorphism and geographical distribution of enKoRV and *phaCin-β*, and compare these to an older related ERV lineage named *phaCin-β-like*, which began colonizing the koala genome between 2.4 and 5.5 million years ago^9^. It is relevant to evaluate the population-wide distribution of these lineages for insights into their infection history, establishment as ERVs and impact on the host population. This is pertinent given the overlapping time intervals of KoRV and *phaCin-β* infections and their potentially continuing activity in the koala population^9^.

For these studies, we take advantage of the Koala Genome Survey^10^, which has produced short-read whole-genome sequences from 430 individual koalas across wild populations in Queensland (QLD; *n*=100), New South Wales (NSW; *n*=246), and Victoria (VIC; *n*=72) as well as two captive populations (*n*=12). We find contrasting polymorphism patterns for the recent ERV lineages across the koala population and, in total, reveal thousands of ERVs, suggesting high ERV burden in koala, especially regarding the two, possibly still active, *phaCin-β* and KoRV lineages. These ERVs reflect historic retrovirus-host interactions and supply markers for improved population conservation genetics.

## Results

### ERV polymorphism

Following our previously described ERV mapping strategy^9,11^ and utilizing reference sequences derived from the three focal ERV lineages KoRV, *phaCin-β*, and *phaCin-β-like*, we analyzed whole-genome short-read sequences from the koala population (Fig. 1a), and identified a total of 12,990 ERV loci. We identified more unique ERV loci in the younger lineages, such that 9,346 loci were classified as KoRV, 3,175 as *phaCin-*β and only 469 as *phaCin-*β*-like* (Table 1). The timing of retroviral activity was also reflected in the ERV frequency trends across the population. The number of individuals having a given KoRV locus varied from 1 to 163 (mean frequency 0.7 %; median frequency 0.5 %). 4,642 (49.7 %) KoRV loci were private, i.e. only found in one individual koala. Individuals had between 2 and 138 KoRV loci in the genome. Population counts per *phaCin-β* locus varied from 1 to 430 (mean frequency 2.2 %; median frequency 0.7 %). Individuals carried 38 to 110 *phaCin-β* loci each, and 1,015 (32.0 %) *phaCin-β* loci were private to a single individual. In contrast, *phaCin-β-like* loci showed higher frequencies (mean frequency 12.9 %; median frequency 1.4 %), reflecting their earlier expansion history, and individuals contained between 27 to 73 *phaCin-β-like* integrations in their genome. There were still relatively many private loci at *phaCin-β-like*, with 151 (32.2 %) loci found in only one individual.

**Table 1.** ERV polymorphism across koala samples (n=430).

| ERV | Total ERVs<sup>1</sup> | Private ERVs | Frequency |  |  | ERVs per koala |
| --- | --- | --- | --- | --- | --- | --- |
|  |  |  | Max | Mean | Median |  |
| KoRV | 9,346 | 4,642 (49.7 %) | 38 % | 0.7 % | 0.5 % | 2 – 138 |
| *phaCin-β* | 3,175 | 1,015 (32.0 %) | 100 % | 2.2 % | 0.7 % | 38 – 110 |
| *phaCin-β-like* | 469 | 151 (32.2 %) | 100 % | 12.0 % | 1.4 % | 27 – 73 |

1. Number of identified locations containing an ERV in at least one individual

**Fig. 1.**
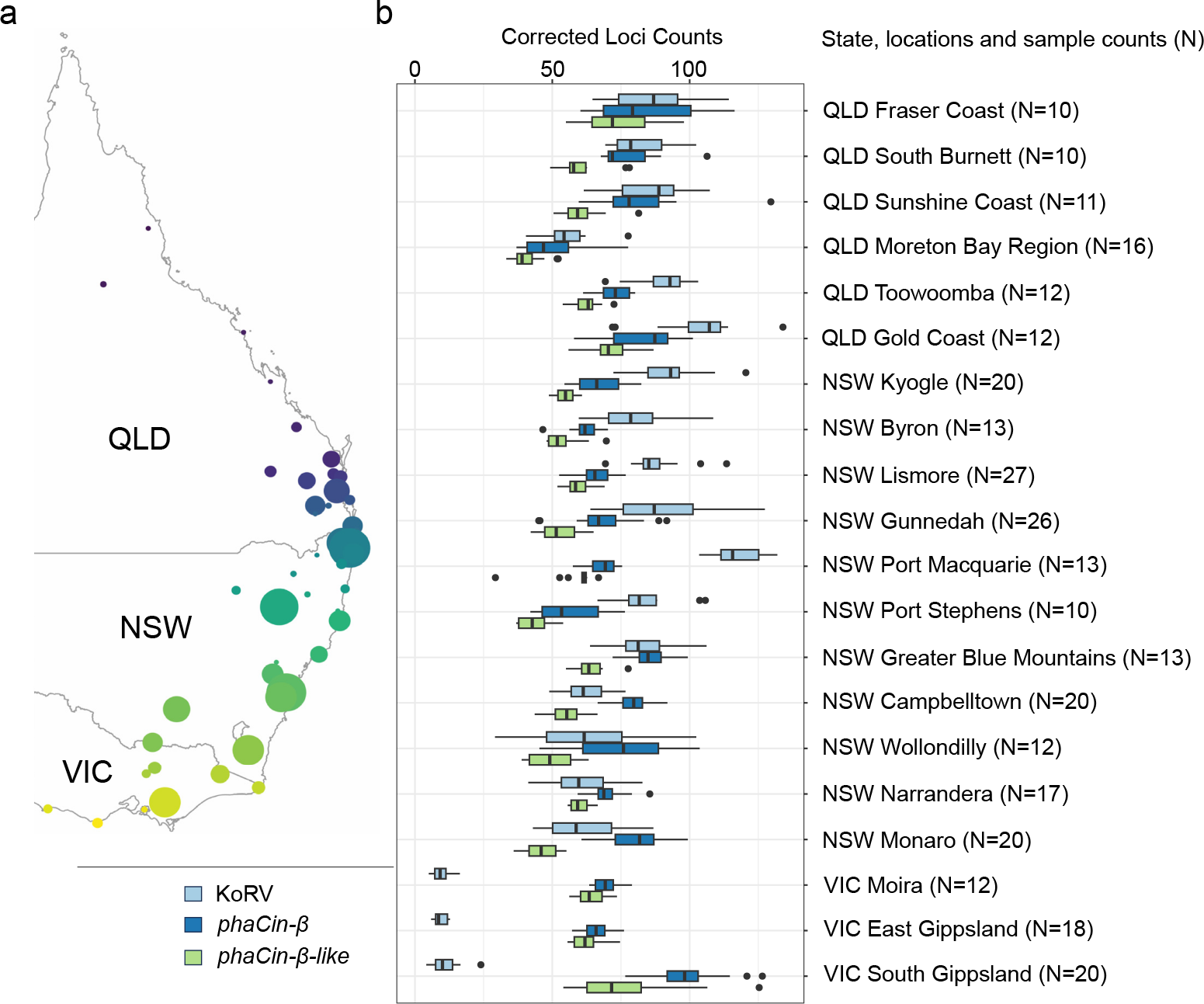
Contrasting ERVs accumulation pattern of across the koala population. **a**, Map of Australian east coast showing koala sampling locations (modified from Hogg et al.^10^). **b**, Boxplots of corrected ERV counts per individual across the sampling locations with more than 10 samples (312 individuals from 20 locations; see details in Supplementary Table 1) for KoRV (light blue), *phaCin-β* (dark blue) and *phaCin-β-like* (green) ERVs. Locations are ordered by latitude.

The identified ERV loci were widely and unevenly distributed across the genome (Supplementary Fig. 1), with indications of integration “hotspots” regions emerging from population wide identifications, for example, a 20 kb region MSTS01000019.1:3,020,001-3,040,000 that contained five *phaCin-β* and five KoRV insertions (Supplementary Fig. 1). This particular region overlaps with a predicted LRP1B gene (MSTS01000019.1:1,511,885-3,684,244), a member of the low-density lipoprotein (LDL) receptor family involved in various normal cell functions and development, and of which disruption is associated with several types of cancer. In some cases, *phaCin-β* and KoRV integrations are particularly close, for example, at MSTS01000042.1:9,858,795-9,858,778 there is only 17 bp separating integrations by *phaCin-β* and KoRV in different samples (Supplementary Fig. 2). These observations support overlapping time of activity for *phaCin-β* and KoRV, since they co-localize to DNA regions accessible at the time of integration. We also observe apparent “coldspot” regions of the genome, such as the 3.7 Mb region MSTS01000019.1:5,903,013-9,671,599 where no integrations by the three focal ERV lineages KoRV, *phaCin-β*, and *phaCin-β-like* were detected in our population-wide screening.

### Geographical distribution of ERV polymorphism

Relative numbers of KoRV, *phaCin-β* and *phaCin-β-like* loci varied across the sampling distribution, most noticeably the difference in KoRV counts between QLD/NSW and VIC (Fig. 1b), reflecting well documented differences in disease prevalence^5,12,13^. Local variation within northern populations is also apparent, such as Port Macquarie samples having comparatively high KoRV counts per individual. In QLD and NSW, the number of KoRV integrations per individual are either comparable or greater than the number of *phaCin-β* integrations. In VIC, the greatest number of integrations per individual were *phaCin-β*, then *phaCin-β-like*, with very few KoRV. Given the stationary lifestyle of koalas, these patterns can reasonably be assumed to reflect the historical intensity of infection, which can thus be inferred to differ significantly between the two retrovirus lineages.

Different patterns of polymorphism and per-locus frequency were observed across the koala distribution (Fig. 2). Frequency comparisons (Fig. 2b,e,h) for NSW/QLD vs VIC for the older ERV lineage, *phaCin-β-like* (Fig. 2b), show strong positive correlation (*r*^2^ = 89.7%), such that integrations at a higher frequency in NSW and QLD are also at higher frequency in VIC. This pattern is largely absent from *phaCin-β* (*r*^2^ = 26.3%) and KoRV (*r*^2^ = 2.1%) (Fig. 2e,h), where intermediate-to-high frequency loci in NSW/QLD are at low frequency, or not present, in VIC, and *vice versa*. These patterns provide strong evidence that the majority of loci in the two younger lineages have occurred after the current koala populations were established, while *phaCin-β-like* loci must pre-date this establishment. Interestingly, there is a small number of *phaCin-β* loci with similar, and high frequencies in both groups, indicating that the infection-window had opened up before the populations fully separated. Pairwise counts of shared integrations (Fig. 2c,f,i) show that there is generally little sharing across wider regions in QLD and NSW regarding *phaCin-β* and KoRV, although some regions show greater locus sharing of both ERV lineages. These regions with greater ERV sharing are likely the result of inbreeding in small, insular populations. Interestingly, there is a large degree of *phaCin-β* locus sharing across VIC, which could be the result of founder effect in this population after reintroduction that is also reflected in the intermediate frequencies of many private *phaCin-β* ERV loci in this state (Supplementary Fig. 3), The *phaCin-β-like* integrations show no apparent pairwise structure across the koala population, consistent with a longer residence in the koala germline.

**Fig. 2.**
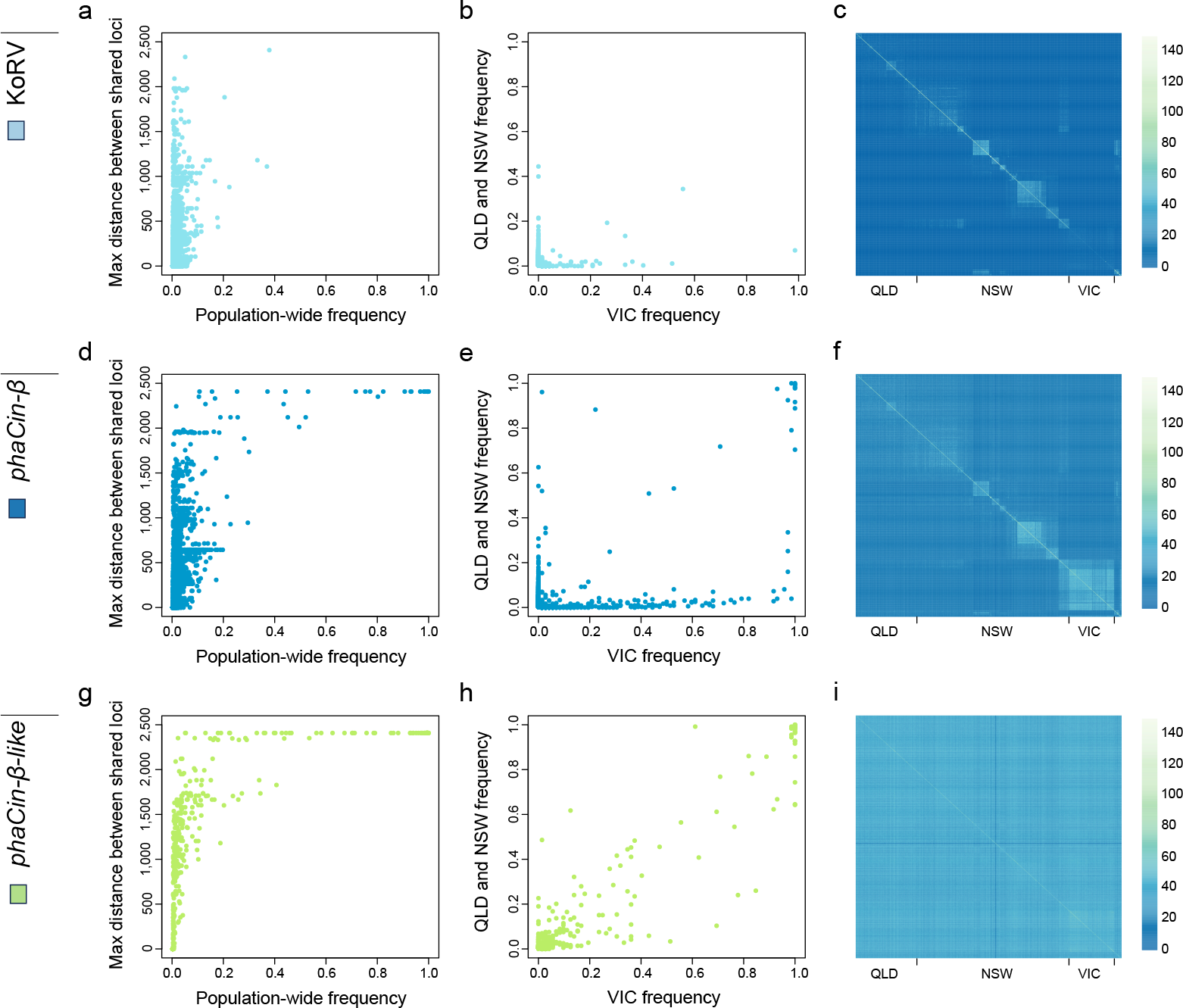
Contrasting geographic ERV frequencies among koalas. Comparisons are shown for the three focal KoRV (**a-c**), *phaCin-β* (**d-f**), and *phaCin-β-like* (**g-i**) ERV lineages across the koala population. **a, d, g**, Maximum geographical (haversine) distance between individuals sharing an ERV integration versus the population-wide frequency for KoRV (**a**), *phaCin-β* (**d**), and *phaCin-β-like* loci (**g**). **b, e, h**, Integration frequency in Queensland (QLD) and New South Wales (NSW) versus frequency in Victoria (VIC) for KoRV (**b**), *phaCin-β* (**e**), and *phaCin-β-like* loci (**h**). **c**,**f**,**i**, Heatmaps of pairwise shared ERV counts between individuals for KoRV (**c**), *phaCin-β* (**f**), and *phaCin-β-like* loci (**i**). Individuals are ordered in sample regions approximately by latitude.

### Comparison of *phaCin-β* insertion polymorphism in koala genomes assemblies

Recently, a second high-quality koala genome assembly became available^14^. The newly sequenced individual was from the bottlenecked South Australian (SA) population, allowing us to compare its *phaCin-β* ERV integrations to those of the reference koala assembly (“Bilbo”), which was from QLD. Using RetroTector^15^ and BLAT^16^, we identified 45 *phaCin-β* integration sites in Bilbo, out of which 17 are solo LTRs and 28 proviruses, ranging between 6,377 and 8,946 bp (median: 8,433 bp). In comparison, the SA genome contains 81 *phaCin-β* integration sites, including 17 solo LTRs and 66 proviruses ranging between 5,638 and 8,518 bp (median: 8,464 bp). A total of 14 integration sites with proviruses and 13 solo LTRs were shared between the two reference genomes (Supplementary Table 2). Of the 28 provirus loci in Bilbo, two (7 %) were solo LTRs and 12 (43 %) were empty pre-integration sites in SA (Supplementary Table 2). Of the 66 provirus loci in the SA assembly, two were solo LTRs (3 %) and 43 (65 %) were empty pre-integration sites in Bilbo (Supplementary Table 2). Like the per-locus frequency differences shown above, the higher number of provirus integrations identified in the SA genome indicates more recent *phaCin-β* activity in the southern koala populations, reflecting a contrasting pattern of historical retrovirus spread to that of KoRV.

## Discussion

Considering the recent listing of the northern koala populations as an endangered species by the Australian Government^17^, it is relevant to improve the population conservation genetics and to better understand the potential threats presented by deleterious genetic elements. Here, we leverage our ERV screening approach^9,11^ across a unique koala population genomics dataset^10^ to compare the polymorphism and distribution pattern of three recent ERV lineages, the *gammaretroviral* KoRV and the two *betaretroviral phaCin-β* and *phaCin-β-like*^9^. The main outcomes from studying these three ERV lineages in population-wide screening are two-fold. First, the observed frequency differences of the three focal ERV lineages across the koala population provide increased resolution over conventional *de novo* single nucleotide polymorphisms (SNPs), due to the abundance of loci segregating at divergent frequencies in different koala populations. This supports accurate tracking of koalas from the different subpopulations for improved conservation genetics. Second, contrasting frequency patterns inform about retroviral infection history of KoRV that is most common in northern koalas and *phaCin-β* found at high frequencies in the south, compared to *phaCin-β-like*, which is a young lineage compared to other ERVs but much older than both KoRV and *phaCin-β*^9^, showing more uniform frequency across the koala population.

We detect thousands of ERV integrations across the koala genome, and decreasing abundance with predicted ERV lineage ages. KoRV, which is the youngest lineage in the study, is the most prolific (*n*=9,346) with high frequency of loci private to individual koalas (49.7 %), reflecting the present KoRV activity in the koala population^3^. Consequently, many of these ERVs are likely to be lost from the population within a few generations. There are fewer than half as many loci of the slightly older *phaCin-β* (*n*=3,175) lineage, with lower frequency of private loci (32%). This lower number, relative to KoRV, could be partly the result of many ERVs being lost due to drift since *phaCin-β* entered the koala population. Alternatively, given the overlapping age estimates between KoRV and *phaCin-β*^9^, differences in numbers of identified loci could also reflect varying degree of activity in the koala population and, by extension, the amount of ERVs were inherited through the germline. Another potential explanation for these patterns could be differences in *phaCin-β* and KoRV integration efficiency. Since *phaCin-β* is estimated to have entered the koala before KoRV, but have fewer ERVs with population-wide spread, it is conceivable that infections were less permissive to ERV establishment or possibly suppressed by the koala. Along these lines, *phaCin-β-like* being the oldest of the three focal ERV lineages identified only a few hundred (*n*=469) loci.

The distribution across the genome by these three ERV lineages was widespread and uneven, showing apparent integration “hotspots”. In particular, hotspot regions sharing many KoRV and *phaCin-β* integrations indicate that specific regions of the genome have been accessible for hundreds of thousands of years. On the other end, there are apparent “coldspots” in the koala genome, where no integrations by our focal three ERV lineages are detected across the koala population. An explanation for these observations could be the fact that some retroviruses have been observed to exhibit integration preference as a result of interactions between their viral structural proteins with host chromatin and cellular proteins^18^. For example, the *lentiviral* HIV-1 displays targeted integrations within gene-dense genomic regions, predominantly within active transcription units^19,20^. Murine leukemia virus (MLV) and other studied *gammaretroviruses* similarly target gene-dense regions, often in the immediate vicinity of promoters and CpG islands^21-23^. Thus, the observed integration patterns in the koala likely reflect the chromatin status of the genome, with accumulation of integrations in accessible and acceptable regions of the genome and absence in regions that are not. Regions of high integration instances from the three ERV lineages may also indicate regions of the genome that have been consistently accessible to retrovirus integration over millions of years.

The widely recognized, major genetic differentiation between Northern (NSW and QLD) and Southern (VIC and SA) koala populations has directed most population management in the past^13,14,24^. The main contributor to this differentiation is the effective extinction and subsequent reintroduction of koalas into the southern states from relict island populations during the early 20^th^ century^14^. As a result, the southern states have very low genetic diversity on the SNP level^24-26^. We observe this genetic divide between the northern and southern states in our data, with a marked difference in KoRV integration counts across the regions. Additionally, we observe evidence for founder effect in the VIC populations, particularly the *phaCin-β* integrations. VIC has prominent locus sharing across the whole sampled region, which would result from founder effect after populations were reintroduced from island refugia. The relative contributions from founder effect on ERV loci, or the potential ongoing *phaCin-β* activity in these southern population requires further study.

Despite the koala populations of the southern states having low genetic diversity^24-26^, we observe considerable insertional diversity by *phaCin-β*, with a substantial number of loci segregating close to 50% allele frequency. While ERV polymorphism can be valuable in tracing the co-evolutionary history of host species and retroviruses, they can also be viewed as informative genetic markers, especially in case of species where relatively recent retroviral activity overlaps with the establishment of present-day population structure. ERVs could be particularly informative genetic markers in conservation genetics, where many species have reduced genetic variation. As in the koala, due to the recent *phaCin-β* activity, ERV loci by this lineage may show higher diversity than SNPs^25^, and could help disentangle genetic relatedness for population conservation management, breeding plans or translocation strategies.

Our study strongly suggests a high ERV burden in koalas, stemming from both the historical expansion of the *phaCin-β* lineage, and the current accumulation of KoRV integrations within QLD and NSW populations. However, the cumulative impact of these ERVs on the fitness of koala individuals remains to be determined. There is indication of *phaCin-β* expression difference between QLD and SA koalas^26^ (note: *phaCin-β* loci were annotated as “syncytin-like”). Due to sequence similarities between *phaCin-β* loci, however, sequencing reads from *phaCin-β* insertions will often mis-map to other *phaCin-β* loci in the reference genome, making it particularly difficult to determine locus specific expression in this ERV lineage. Nevertheless, expression of *phaCin-β* ERVs detected in SA and QLD imply that they may have a functional impact and that further work is required to determine these potential effects.

In conclusion, we present contrasting polymorphism patterns for three, in evolutionary terms, recent ERV lineages across the koala population and reveal thousands of ERVs among the 430 sequenced koala individuals. We show clear population differences for the most recently, possibly still active, *phaCin-β* and KoRV lineages that illuminate recent retrovirus-host interactions and present a resource for population conservation genetics of the endangered koala.

## Methods

### Whole genome sequencing data

430 koala alignment files (bams; average coverage: 32.3x, range: 11.3–66.8x) were accessed from the Amazon Web Services Open Data platform (https://registry.opendata.aws/australasian-genomics; accessed 2023/04/11)^10^. Visualization and inspection of candidate loci used the Integrative genomics viewer, IGV^27^.

### Polymorphic reference ERV loci

Deletions were detected using Delly v 0.7.7^28^ and Lumpy v 0.3.0^29^, then compared to the koala candidate ERV regions list that was formed from RetroTector ERVs in R^30^ using the package Genomic Ranges^31^, regions identified by BLAT with similarity to full length ERVs in the three focal expansions (non-overlapping with RetroTector identified loci), and regions identified by BLAT with similarity to the LTRs of ERVs the three focal expansions (non-overlapping with the two previous datasets). Inaccurate Delly calls for KoRVs were removed after visual inspection. For deletions called by both softwares with comparable lengths, the Delly genotypes were retained. Unique Delly and Lumpy calls were both kept.

### Non-reference ERV insertional loci

Insertions identified by Retroseq^32^ as previously described^9^. Briefly, ERV loci were detected by RetroSeq^32^ using a custom ERV reference library (-eref), formed from Koala ERVs and retrovirus sequences from diverse lineages. Retroseq *call* using softclips and minimum two reads were used across the samples. Calls were filtered in R to those with FL 8, CLIP3 >= 2 and CLIP5 >= 2, cov/2 <= minGQ <= 2*cov in order to define ERV loci, which was then restricted to the three focal lineages: KoRV, *phaCin-β* and *phaCin-β-like* (using a strict filter that all ERVs called at a locus were all called for the same lineage). During this process, a number of putative Retroseq loci were identified as containing calls for two ERV lineages and were inspected manually. After inspection, 24 such loci that contained two insertions by different lineages separated by only a very small distance were manually curated. Overlaps to reference ERVs were removed unless insertion was identified in another ERV lineage, as this represented secondary insertion. RetroSeq filters were relaxed (FL3; GC2) to count common ERV insertions across individuals.

### Genomic locations of ERV integrations

KoRV, *phaCin-β* and *phaCin-β-like* ERV accumulation in the koala genome was inspected by plotting locus midpoints across the largest scaffolds in the reference assembly. Genomic regions (20 kb windows) with multiple integrations were identified using a binnedSum in R.

### Population comparisons

ERV counts per individual were adjusted to account for differences in coverage (corrcount = ERV count within sample / sample coverage * mean coverage for all samples). Plots were generated using R and ggplot^33^. Heatmaps were generated with the pheatmap package in R.

### Comparisons between reference assemblies

BLAT was used to identify *phaCin-β* loci in the South Australian (SA) koala assembly (Genbank accession: GCA_030178435.1), using both full-length *phaCin-β* and LTR sequences. Flanking regions (1kb upstream and downstream) of candidate proviruses and solo LTRs were extracted from the SA assembly and lifted over to the reference genome using BLAT. The reverse process then lifted over *phaCin-β* loci from the reference genome to the SA assembly. Comparisons between these two assemblies were manually inspected and summarized.

## Data availability

The reference koala assembly is available at Genbank: GCA_002099425.1 [https://www.ncbi.nlm.nih.gov/datasets/genome/GCF_002099425.1/]. The South Australian koala assembly is available at Genbank: GCA_030178435.1 [https://www.ncbi.nlm.nih.gov/datasets/genome/GCA_030178435.1/]. Whole genome sequencing data is available from Amazon Web Services Open Data platform (https://registry.opendata.aws/australasian-genomics/)^10^.

## Code availability

Code and supporting files are available at GitHub: (https://github.com/PatricJernLab/Koala_ERVs_population_screening: DOI PLACEHOLDER).

## Acknowledgements

We would like to thank Jason Hill for helpful suggestions and comments. Data was produced as part of the Koala Genome Survey with funding from the NSW Government and the Australian Government’s Bushfire Recovery for Wildlife and their Habitats program (GA-2000526), further support was provided by The University of Sydney, Amazon Web Services Open Data Sets, Ramaciotti Centre for Genomics and Illumina. The data was derived from samples provided by Amber Gillett, Amy Shima, Australian Museum, Australian Museum Frozen Tissue Collection, Ben Moore, Carolyn Hogg, Enhua Lee, Fiona Hogan, Karen Marsh, Lachlan Wilmott, Lyndal Husle, Mark Krockenberg, Michaela Blyton, Peter Timms, Rachel Labador, and Taronga Western Plains Zoo. This work was funded by Swedish Research Council VR Grants 2021-01740 (to P.J.) and 2021-04238 (to M.L.) and FORMAS Grant 2018-01008 (to P.J.). The computations/data handling were provided by the Swedish National Infrastructure for Computing (SNIC) at the Uppsala Multidisciplinary Center for Advanced Computational Science, partially funded by the Swedish Research Council through Grant Agreement 2018-0597 (Projects SNIC2022/22-945, NAISS-2023/23-14).

## Author Contributions

M.L., M.E.P. and P.J. designed research; M.L. carried out endogenous retrovirus genotyping; M.L., M.E.P., and P.J. analyzed data; M.L. drafted the initial manuscript. M.L. and P.J. wrote the paper.

## Additional information

### Supplementary information

The online version contains supplementary material available at xxxxx.

