## Supplementary appendix for "Contrasting segregation patterns among endogenous retroviruses across the koala population"

*Correspondence:

Supplementary Fig. 1 – 3

Supplementary Table 1 – 2

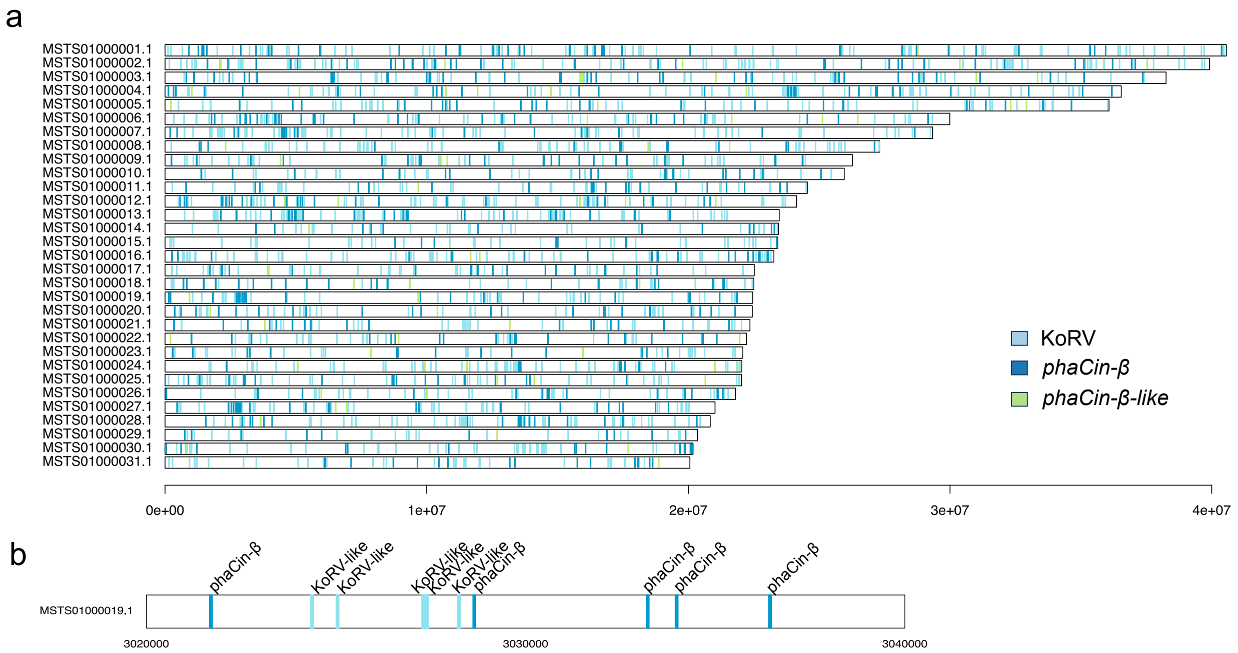


**Supplementary Fig. 1 | Population-wide ERV locations along chromosomes. a,** Positions of mapped ERVs along the largest koala genome scaffolds (assembled lengths >20 MB) with indicated KoRV, *phaCin-β* and *phaCin-β-like* loci. **b,** Example region on koala scaffold MSTS01000019.1:3,020,000-3,040,000 where 10 loci corresponding to *phaCin-β* (n=5) and KoRV (n=5) were identified across the koala population.

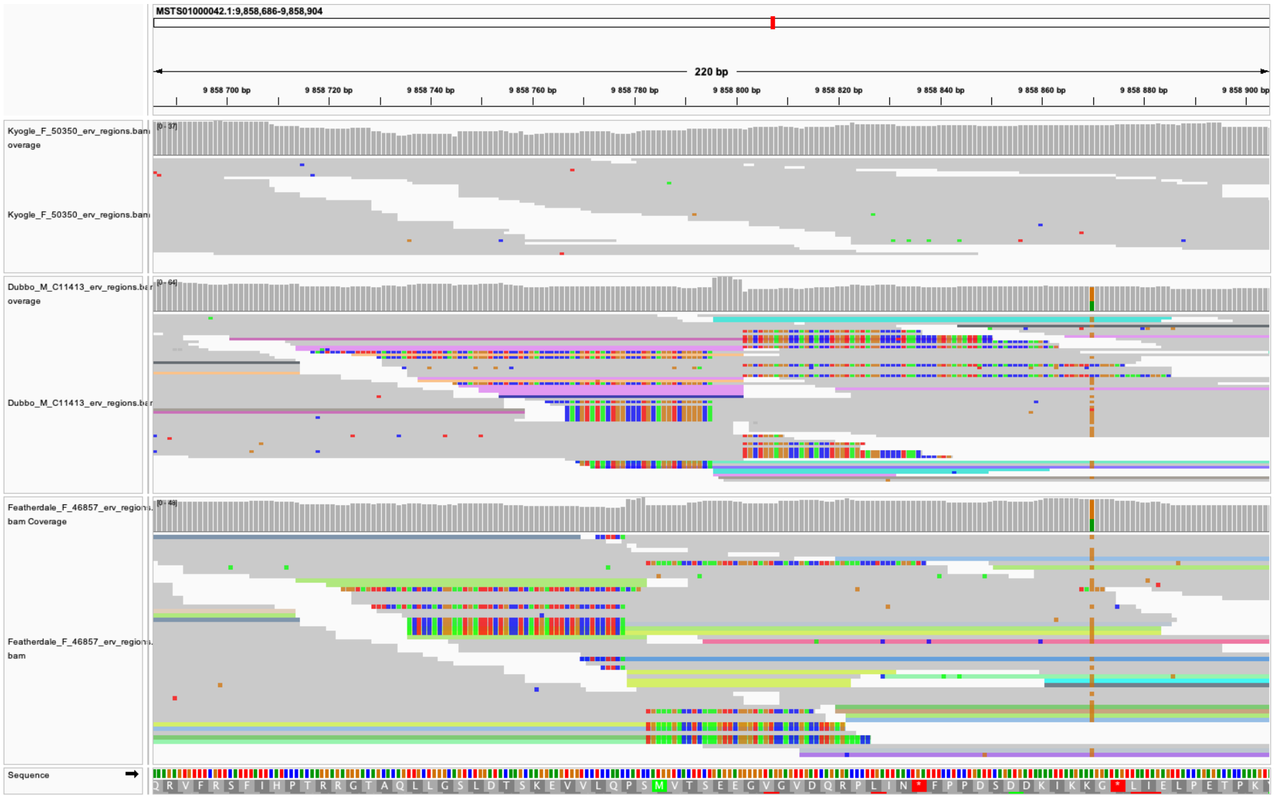


**Supplementary Fig 2. | ERV locus confirmation**. IGV view of koala genomic region MSTS01000042.1:9,858,686-9,858,904, showing ERV loci identified within close genomic proximity in the koala population. The top panel (Kyogle_F_50350) shows no integrations in this range, the middle panel (Dubbo_M_C11413) indicates a *phaCin-β* integration (position 9,858,795, with a 6 bp target site duplication), and the bottom panel (Featherdale_F_46857) has a KoRV integration (position 9,858,778, with a 4 bp target site duplication).

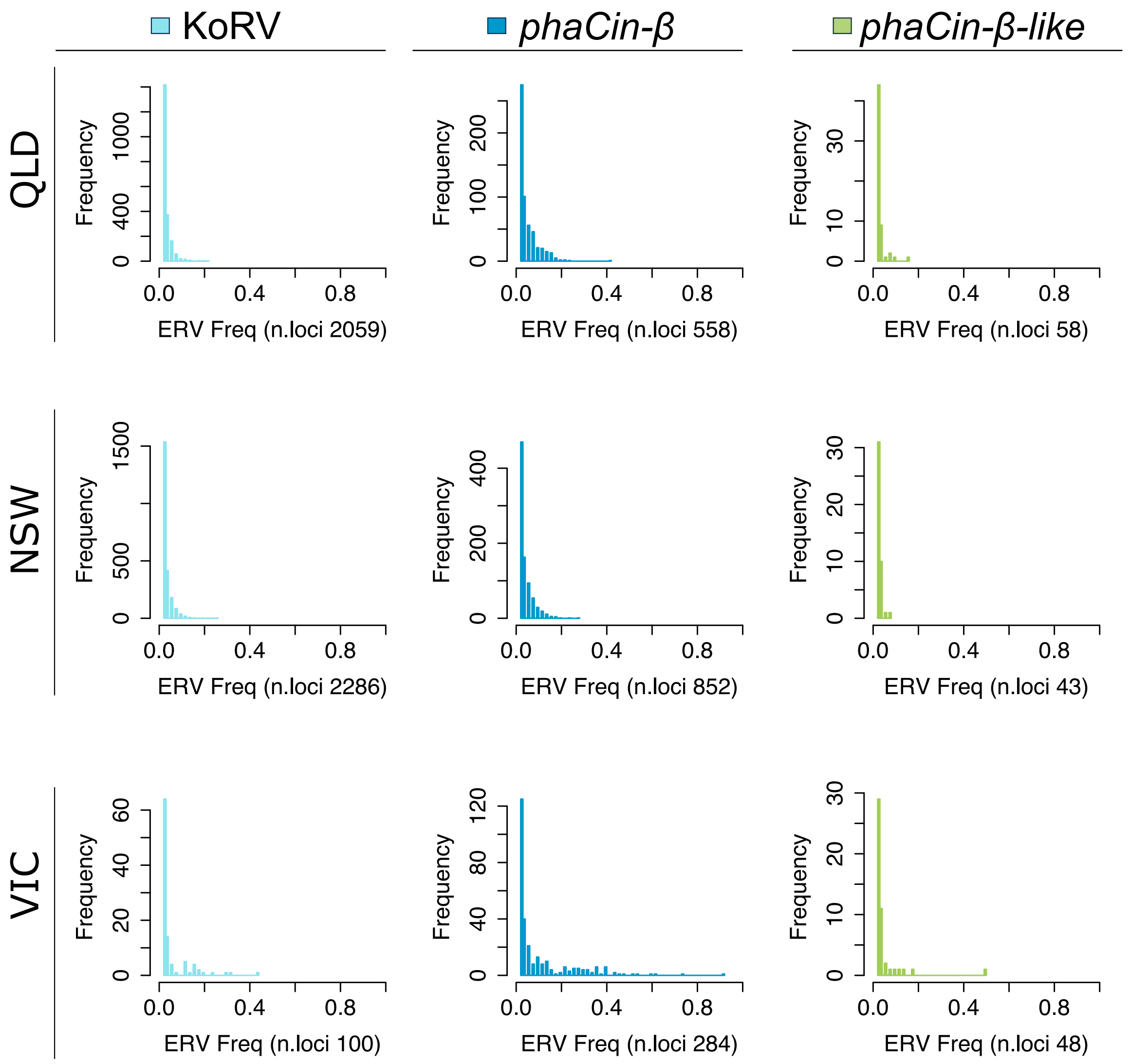


**Supplementary Fig. 3 | Histogram of ERV frequencies across Australian states.** Queensland (QLD; upper row), New South Wales (NSW; middle row) and Victoria (VIC; lower row) for the three ERV lineages KoRV (left column), *phaCin-β* (middle column) and *phaCin-β-like* (right column). From each state, 50 individuals were randomly sampled, private ERVs to the state identified, and the frequencies of these plotted as histograms (number of private ERVs for each state and ERV lineage indicated by "n.loci"). Similar private ERV frequency patterns can be seen in QLD and NSW for both KoRV and *phaCin-β*, however frequencies in VIC show a contrasting pattern, with intermediate to high frequencies in the state.

**Supplementary Table 1 | Sample locations and number of included samples (N) in the koala population ERV screening.**

| Location (Amazon Web Service Folder Name) | Included sampling sites | N |
|---|---|---|
| QLD Fraser Coast | Fraser Coast | 10 |
| QLD South Burnett | South Burnett | 10 |
| QLD Sunshine Coast | Sunshine coast and Noosa | 11 |
| QLD Moreton Bay Region | Moreton Bay | 16 |
| QLD Toowoomba | Toowoomba | 12 |
| QLD Gold Coast | Gold Coast | 12 |
| NSW Kyogle | Kyogle and Northern Rivers | 20 |
| NSW Byron | Byron Byron bay and Northern Rivers | 13 |
| NSW Lismore | Lismore and Northern Rivers | 27 |
| NSW Gunnedah | Gunnedah and Liverpool Plains | 26 |
| NSW Port Macquarie | Port Macquarie | 13 |
| NSW Port Stephens | Port Stephens | 10 |
| NSW Greater Blue Mountains | Blue Mountains | 13 |
| NSW Campbelltown | Campbelltown and SW Sydney | 20 |
| NSW Wollondilly | STHD | 12 |
| NSW Narrandera | Narrandera | 17 |
| NSW Monaro | Monaro | 20 |
| VIC Moira | Murray river | 12 |
| VIC East Gippsland | Gelantipy and Mallacoota | 18 |
| VIC South Gippsland | Gippsland and SHGLD | 20 |

**Supplementary Table 2 | Homologous *phaCin-β* loci structural variations.** Each *phaCin-β* locus is characterized as either pre-integration site (0), provirus (1) or solo-LTR (2) in homologous koala reference assembly (Bilbo) and South Australian assembly (SA) regions.

| Locus Bilbo | Locus SA | Bilbo | SA |
|---|---|---|---|
| NW_018343952.1:16776779-16784633 | JAOEJA010000741.1:3693979-3700585 | 1 | 1 |
| NW_018343953.1:25191880-25191879 | JAOEJA010000383.1:15367455-15367785 | 0 | 2 |
| NW_018343953.1:3488650 | JAOEJA010000383.1:37095927-37101650 | 0 | 1 |
| NW_018343954.1:1113280 | JAOEJA010001327.1:16494524-16502992 | 0 | 1 |
| NW_018343956.1:10725171-10725519 | JAOEJA010000643.1:166112233-166112582 | 2 | 2 |
| NW_018343956.1:31726436 | JAOEJA010000643.1:187134225-187139869 | 0 | 1 |
| NW_018343957.1:19834202-19842624 | JAOEJA010000514.1:127320655-127329107 | 1 | 1 |
| NW_018343960.1:17656158 | JAOEJA010000073.1:8568964-8577431 | 0 | 1 |
| NW_018343961.1:21305097-21312174 | JAOEJA010000656.1:72156087 | 1 | 0 |
| NW_018343962.1:16321441 | JAOEJA010001159.1:68645483-68653953 | 0 | 1 |
| NW_018343962.1:19529052 | JAOEJA010001159.1:71869310-71877720 | 0 | 1 |
| NW_018343968.1:2177802 | JAOEJA010000518.1:69672434-69680900 | 0 | 1 |
| NW_018343969.1:1487319 | JAOEJA010000885.1:1040756-1046403 | 0 | 1 |
| NW_018343970.1:10909918-10918301 | JAOEJA010000399.1:21894572-21903025 | 1 | 1 |
| NW_018343970.1:2821102 | JAOEJA010000399.1:13785669-13794141 | 0 | 1 |
| NW_018343971.1:7508696-7517168 | JAOEJA010000115.1:48719805 | 1 | 0 |
| NW_018343972.1:17325482-17333901 | JAOEJA010000594.1:17459762-17468193 | 1 | 1 |
| NW_018343972.1:19061308 | JAOEJA010000594.1:19196101-19204562 | 0 | 1 |
| NW_018343977.1:8895922-8896230 | JAOEJA010000643.1:131893467-131899831 | 2 | 1 |
| NW_018343978.1:1004264-1004615 | JAOEJA010000937.1:38830715-38831067 | 2 | 2 |
| NW_018343978.1:17507321-17515793 | JAOEJA010000937.1:22311387-22311381 | 1 | 0 |
| NW_018343979.1:14125670-14134137 | JAOEJA010000643.1:107090776-107090770 | 1 | 0 |
| NW_018343981.1:16882197-16890704 | JAOEJA010000824.1:18052340-18052333 | 1 | 0 |
| NW_018343981.1:20154753 | JAOEJA010000824.1:21320126-21320474 | 0 | 2 |
| NW_018343981.1:913805-922217 | JAOEJA010000097.1:7443712-7452168 | 1 | 1 |
| NW_018343986.1:17531244-17539561 | JAOEJA010000701.1:16720294-16720287 | 1 | 0 |
| NW_018343987.1:24805-33210 | JAOEJA010000073.1:88742554-88751006 | 1 | 1 |
| NW_018343990.1:17700228-17707162 | JAOEJA010000575.1:15013365-15020311 | 1 | 1 |
| NW_018343991.1:9188833 | JAOEJA010000639.1:21949905-21958377 | 0 | 1 |
| NW_018343992.1:15851813 | JAOEJA010000399.1:60731282-60739766 | 0 | 1 |
| NW_018343998.1:13200036 | JAOEJA010000936.1:26338724-26347182 | 0 | 1 |
| NW_018344015.1:11021055 | JAOEJA010000594.1:33520830-33529288 | 0 | 1 |
| NW_018344016.1:11645789-11654227 | JAOEJA010000185.1:57612826-57613147 | 1 | 2 |
| NW_018344023.1:1970208-1970556 | JAOEJA010000042.1:8506053 | 2 | 0 |
| NW_018344027.1:5576604-5585072 | JAOEJA010001159.1:90961322 | 1 | 0 |
| NW_018344028.1:10202984-10211399 | JAOEJA010000932.1:10200956-10209393 | 1 | 1 |
| NW_018344031.1:1267997-1276433 | JAOEJA010000468.1:42278665-42287131 | 1 | 1 |
| NW_018344031.1:4359211 | JAOEJA010000468.1:39197815-39206277 | 0 | 1 |
| NW_018344032.1:61864 | JAOEJA010000795.1:33319-38959 | 0 | 1 |

*Continued.*

| Locus Bilbo | Locus SA | Bilbo | SA |
|---|---|---|---|
| NW_018344034.1:1625624 | JAOEJA010000922.1:23659367-23665004 | 0 | 1 |
| NW_018344035.1:3747592 | JAOEJA010000825.1:40756735-40765207 | 0 | 1 |
| NW_018344036.1:5232965-5241393 | JAOEJA010000643.1:24211526-24219993 | 1 | 1 |
| NW_018344038.1:2683935-2684249 | JAOEJA010000073.1:125781336-125781651 | 2 | 2 |
| NW_018344038.1:5078926-5085302 | JAOEJA010000073.1:128190050-128196418 | 1 | 1 |
| NW_018344038.1:6202006-6202340 | JAOEJA010000073.1:129318884-129319219 | 2 | 2 |
| NW_018344042.1:4319930-4328398 | JAOEJA010000659.1:6771828 | 1 | 0 |
| NW_018344043.1:6379136 | JAOEJA010000112.1:6159309-6167774 | 0 | 1 |
| NW_018344044.1:9997139 | JAOEJA010000548.1:1081107-1089573 | 0 | 1 |
| NW_018344046.1:1636918-1645388 | JAOEJA010001082.1:40532856 | 1 | 0 |
| NW_018344050.1:2061073-2061422 | JAOEJA010000656.1:50143272-50143623 | 2 | 2 |
| NW_018344052.1:5639248-5639578 | JAOEJA010000944.1:5007950 | 2 | 0 |
| NW_018344055.1:3287799 | JAOEJA010000514.1:10616371-10624381 | 0 | 1 |
| NW_018344060.1:1883262-1891724 | JAOEJA010000050.1:49846749 | 1 | 0 |
| NW_018344064.1:5611633-5611961 | AOEJA010000001.1:6481120-6481449 | 2 | 2 |
| NW_018344073.1:88015 | JAOEJA010000001.1:10688970-10697435 | 0 | 1 |
| NW_018344077.1:413575-422011 | JAOEJA010000825.1:8970417-8978909 | 1 | 1 |
| NW_018344079.1:2092946-2101182 | JAOEJA010000643.1:79752382-79760825 | 1 | 1 |
| NW_018344081.1:5419770 | JAOEJA010000660.1:7293343-7301812 | 0 | 1 |
| NW_018344088.1:306475 | JAOEJA010000399.1:2812629-2821097 | 0 | 1 |
| NW_018344090.1:7142871 | JAOEJA010000077.1:7243472-7251935 | 0 | 1 |
| NW_018344092.1:5237814-5246236 | JAOEJA010000937.1:45088459 | 1 | 0 |
| NW_018344094.1:6448134 | JAOEJA010000655.1:13805733-13814250 | 0 | 1 |
| NW_018344101.1:3685364-3685714 | JAOEJA010000828.1:3826629-3826978 | 2 | 2 |
| NW_018344115.1:5979649 | JAOEJA010000383.1:50945951-50954417 | 0 | 1 |
| NW_018344118.1:470197-470545 | JAOEJA010000884.1:24680041-24680389 | 2 | 2 |
| NW_018344126.1:3147639 | JAOEJA010001090.1:23275883-23284345 | 0 | 1 |
| NW_018344136.1:2866032 | JAOEJA010000936.1:19877661-19886129 | 0 | 1 |
| NW_018344141.1:2337383 | JAOEJA010001160.1:2328151-2336620 | 0 | 1 |
| NW_018344141.1:3743958-3744286 | JAOEJA010001160.1:3743056-3743385 | 2 | 2 |
| NW_018344142.1:3232771 | JAOEJA010000185.1:3362695-3371163 | 0 | 1 |
| NW_018344150.1:5531959 | JAOEJA010000279.1:4281625-4290094 | 0 | 1 |
| NW_018344160.1:4450675 | JAOEJA010000824.1:1026382-1034849 | 0 | 1 |
| NW_018344170.1:3741391 | JAOEJA010000518.1:91250655-91256307 | 0 | 1 |
| NW_018344175.1:3999554 | JAOEJA010000073.1:84691974-84692323 | 0 | 2 |
| NW_018344182.1:3982440-3982767 | JAOEJA010001333.1:6136880 | 2 | 0 |
| NW_018344205.1:202785 | JAOEJA010001284.1:3547825-3556289 | 0 | 1 |
| NW_018344206.1:2610228-2610577 | JAOEJA010000932.1:15168892-15169242 | 2 | 2 |
| NW_018344231.1:3086220 | JAOEJA010000073.1:57018669-57027136 | 0 | 1 |
| NW_018344236.1:145019-145341 | JAOEJA010000279.1:14361675-14370112 | 2 | 1 |
| NW_018344242.1:2231815-2232144 | JAOEJA010001095.1:16710732-16711062 | 2 | 2 |
| NW_018344243.1:692853-701323 | JAOEJA010000050.1:75477209 | 1 | 0 |
| NW_018344260.1:682177 | JAOEJA010000548.1:26237512-26245975 | 0 | 1 |
| NW_018344287.1:523019-523013 | JAOEJA010000531.1:589831-598299 | 0 | 1 |
| NW_018344297.1:1640839-1649312 | JAOEJA010000941.1:751658-760157 | 1 | 1 |

*Continued.*

| Locus Bilbo | Locus SA | Bilbo | SA |
|---|---|---|---|
| NW_018344347.1:460174 | JAOEJA010000279.1:562112-570579 | 0 | 1 |
| NW_018344443.1:40288-40637 | JAOEJA010000270.1:104856-105205 | 2 | 2 |
| NW_018344501.1:34883-43828 | JAOEJA010000138.1:475230-475698 | 1 | 2 |
| NW_018344659.1:26558 | JAOEJA010000275.1:88793-97260 | 0 | 1 |
| NW_018345530.1:12850 | JAOEJA010000497.1:52250-60744 | 0 | 1 |
| NW_018345562.1:10256 | JAOEJA010000312.1:335038-343506 | 0 | 1 |
| NW_018345562.1:6426 | JAOEJA010001139.1:68502-76969 | 0 | 1 |
| NA | JAOEJA010000872.1:5066-13211 | NA | 1 |
| NA | JAOEJA010001074.1:18549-27019 | NA | 1 |
| NA | JAOEJA010000062.1:43644-52104 | NA | 1 |
| NA | JAOEJA010000192.1:0-8461 | NA | 1 |
| NA | JAOEJA010000262.1:14610-20253 | NA | 1 |
| NA | JAOEJA010000604.1:8678-17053 | NA | 1 |
| NA | JAOEJA010000871.1:5150-13598 | NA | 1 |
